# From recognition to neglect: Molecular and physiological responses to heterospecific pollen decay with evolutionary distance

**DOI:** 10.64898/2026.06.02.729600

**Authors:** Rachel O. Cohen, Alessandra Masi, Sarah M. Chin, Joseph H. Williams, Deren A.R. Eaton

## Abstract

Post-mating, pre-zygotic (PMPZ) reproductive barriers are often attributed to precise molecular recognition between male and female gametes, yet little is known about how these interactions change as species diverge. In flowering plants, pollen tube growth depends on coordinated signaling between pollen and pistil, raising the question of whether PMPZ barriers arise through active incompatibility mechanisms or gradual loss of pollen–pistil coordination. Here, we combined transcriptomic profiling and pollen tube growth assays across a phylogenetically structured set of crosses in a diverse alpine plant community. We show that the magnitude of the pistillar transcriptomic response to pollination declined quantitatively with evolutionary distance rather than shifting in a binary compatible/incompatible manner. Pollination-associated functional gene ontology categories were strongest in conspecific and intrageneric crosses and weakened with divergence. Heterospecific pollen tubes also grew more slowly than conspecific tubes, with growth rates declining overall with genetic distance. However, the slowest growth occurred in intrageneric crosses, suggesting that close relatives may represent a distinct evolutionary zone where reduced maternal support coincides with additional hindrance mechanisms. Together, these results support a graded, divergence-dependent model of pollen–pistil incongruence driven primarily by attenuation of coordinated growth rather than strict heterospecific rejection.

## Introduction

Reproductive success in sexually reproducing organisms depends on coordinated interactions between male and female tissues. Across multicellular eukaryotes, these interactions continue well after mating (*i.e.* gamete transfer), unfolding predominantly within female reproductive environments that can influence whether and how fertilization proceeds. Post-mating, pre-zygotic (PMPZ) processes occur after male gametes (or male gametophytes) make contact with female reproductive tissues and prior to fertilization and zygote formation. Because these processes are mediated by molecular recognition, cellular signaling, and physiological responses that shape the performance of male gametes within female tissues (Swanson and Vacquier 2002), they have been widely discussed as key contributors to reproductive isolation and speciation (Howard 1999; Coyne and Orr 2004; Garlovsky et al. 2024). However, PMPZ processes also have direct consequences for female reproductive fitness: male gametes from other species can alter resource allocation, disrupt the performance of conspecific gametes, or impose physiological costs on female tissues, even when hybrid formation never occurs (Ashman and Arceo-Gómez 2013; Ting et al. 2014; Garlovsky and Snook 2018; Larson et al. 2019). Despite their importance, PMPZ barriers remain the least studied class of reproductive barriers (Garlovsky et al. 2024), and we know relatively little about how recognition systems within female reproductive tissues detect and regulate the performance of male gametes originating from different species.

In many systems, the very fact that fertilization is an internal process has hampered the study of PMPZ processes. Flowering plants, though, provide a powerful system for examining these interactions. Whereas motile sperm may be difficult to localize in animal systems, the growth of pollen tubes occurs within accessible maternal tissues and depends on sustained coordination between male and female partners. Following deposition on a receptive stigma, compatible pollen is hydrated, then germinates a pollen tube which penetrates the stigmatic surface and elongates through the style. Although pollen tubes can grow in the absence of maternal tissues, elongation is typically slower and less efficient *in vitro*, indicating that active support from the pistil enhances pollen tube performance *in vivo* (Sanders and Lord 1992; Herrero and Hormaza 1996; Qin et al. 2009). Successful conspecific PMPZ interactions involve coordinated chemical, physiological, and transcriptional responses in both pollen and pistil, including modulation of ion fluxes and reactive oxygen species, cell wall remodeling, cytoskeletal dynamics, and peptide- and receptor-mediated signaling that guide and sustain polarized tip growth (Krichevsky et al. 2007; Bedinger et al. 2017; Mizuta and Higashiyama 2018). Hundreds of genes contribute to pollen-pistil interactions, forming an integrated molecular network that supports pollen tube elongation *in vivo*.

When pollen and pistil originate from different species, this coordination can fail. Such breakdowns in compatibility have been described as “incongruence,” reflecting insufficient matching between interacting male and female components to sustain normal pollen tube performance (de Nettancourt 1997; Hogenboom et al. 1997; Bedinger et al. 2017). Although the genes and pathways underlying successful conspecific growth are increasingly well-characterized, less is known about how these networks respond to heterospecific pollen. For example, it remains unclear whether genetic and physiological responses break down abruptly or decay gradually with evolutionary divergence, or whether particular pathways consistently fail before others. Thus, while incongruence is often observed phenotypically as reduced pollen tube growth, we know little of the molecular mechanisms that may be responsible for this breakdown in coordination. Whether the mechanisms responsible for pollen-pistil incongruence are shared across lineages (*e.g.* through convergent or parallel evolution) or are lineage specific remains a major outstanding question.

In natural plant communities, opportunities for heterospecific pollen receipt are common. Many species flower synchronously and share pollinators, resulting in frequent interspecific pollen exchange even among taxa that rarely hybridize (Kohn and Waser 1985; Ashman and Arceo-Gómez 2013; Moreira-Hernández and Muchhala 2019). Experimental manipulations show that increasing the proportion of heterospecific pollen can reduce seedset, often in proportion to its abundance (Ashman and Arceo-Gómez 2013; Briggs et al. 2016; Moreira-Hernández and Muchhala 2019). Although this heterospecific pollen transfer negatively impacts both male and female reproductive success through pollen loss and seedset reductions, respectively, disruption of mating via heterospecific pollen transfer will be more costly to the maternal parent because of greater energetic investment in each gamete. Maternal regulation of heterospecific pollen tube growth could mitigate the fitness costs imposed by heterospecific pollen receipt, and, because of these higher fitness costs in the maternal parent, maternal tissues are more likely to evolve mechanisms that discriminate against heterospecific pollen.

The expectation that pistil-acting mechanisms actively suspend heterospecific pollen germination and pollen tube growth has been hypothesized in many systems (Ashman and Arceo Gomez 2013, 2016; Moreira-Hernandez and Muchhala 2019). Evidence for pistil-acting mechanisms that disrupt heterospecific pollen performance, though, remains mixed. Most evidence for active rejection of pollen tubes in the flowering plants comes from self-incompatibility (SI) systems, which provide clear examples of pistil-acting genes functioning in the recognition and rejection of self pollen (Takayama and Isogai 2005). Indeed, expansive work on SI systems is likely the basis for the expectation that pistils are capable of suspending heterospecific pollen (Nasrallah and Nasrallah 1993; Hormaza and Herrero 1994; Hirose et al. 1995; Luu et al. 2000; Richman and Kohn 2000; Takayama and Isogai 2005). There is some evidence that SI mechanisms may be recruited for the rejection of heterospecific pollen; however, the genes underlying SI-mediated rejection have been studied primarily in the context of intraspecific crosses. Moreover, when SI systems are implicated in interspecific reproductive barriers, they are most often discussed in relation to asymmetries in compatibility under the SI × SC rule (Baek et al. 2015; Bedinger et al. 2017; Broz and Bedinger 2021). Nevertheless, because heterospecific pollen receipt can impose substantial fitness costs, active recognition and rejection of heterospecific pollen by pistil-acting mechanisms—whether derived from SI systems or other pathways—remains a leading hypothesis for the origin of pollen–pistil incongruence (Ashman and Arceo-Gómez 2013; Bedinger et al. 2017; Arceo-Gómez et al. 2019; Moreira-Hernández and Muchhala 2019; Broz and Bedinger 2021).

Recently, Cao et al. (2025) demonstrated a molecular mechanism distinct from SI by which species in Brassicaceae actively suspend heterospecific pollen germination, providing direct evidence of targeted interspecific rejection. However, such mechanisms appear to be lineage-specific, and, being outside classical incompatibility systems, may involve many genes of minor effect, making them more difficult to discover. Some studies report that pollen from closely related species performs better than that from more distant relatives (Streher et al. 2020), whereas others find no relationship between phylogenetic distance and heterospecific pollen tube germination or growth (Cohen et al. 2025b). To our knowledge, no study has quantified pistillar transcriptional responses to heterospecific pollination across a phylogenetically structured gradient of donor relatedness.

Besides maternal rejection of heterospecific pollen, an additional possibility is that pistils preferentially support conspecific or pollen from closely related species. Because pollen tubes are capable of elongation in the absence of maternal tissues, differences in performance may reflect variation in maternal enhancement rather than active inhibition of heterospecific pollen tubes (Sanders and Lord 1989; Herrero and Hormaza 1996; Lin et al. 2014; Losada and Herrero 2014). Pistils provide chemical guidance cues and structural environments that facilitate sustained polarized growth (Krichevsky et al. 2007; Lora et al. 2016; Mizuta and Higashiyama 2018; Riglet et al. 2020), and coordinated changes in gene expression have been shown to accompany successful conspecific interactions (Bedinger et al. 2017). Under this framework, we expect a progressive loss of matched signaling and physiological integration as genetic divergence increases, rather than discrete incompatibility responses.

Here, we investigate how heterospecific pollen-pistil interactions vary across evolutionary distance within a diverse alpine plant community. Using controlled pollinations, we compare conspecific, closely related heterospecific, and distantly related heterospecific crosses. We sequence RNA from pistils following pollination to characterize transcriptional responses to pollen identity, and measure pollen tube growth rates in parallel to assess physiological consequences. This design allowed us to test whether pistillar gene expression responses track phylogenetic distance, and if variation in molecular response predicts differences in pollen tube performance.

## Methods

### Study system

This study was conducted in subalpine meadows surrounding Rocky Mountain Biological Laboratory (RMBL) in Gothic, Colorado, USA, where many plant species flower synchronously and share generalist pollinators. We focused on two focal maternal species, *Pedicularis groenlandica* (Orobanchaceae) and *Mimulus guttatus* (Phrymaceae), and a set of closely and distantly related taxa that co-occur in their community. *P. groenlandica* is a facultative hemiparasite with well-characterized pollination biology (Macior 1970, 1983; Eaton et al. 2012; Tong and Huang 2016; Liang et al. 2018; Cohen et al. 2025b). *Mimulus guttatus* is a model system in plant ecology and evolution, with extensive work on hybridization and reproductive barriers (Ramsey et al. 2003; Wu et al. 2008; Coughlan et al. 2020; Nelson et al. 2021). Chromosome-scale reference genomes are available for both species and were used for transcriptomic analyses. *P. groenlandica* served as maternal recipients in both transcriptomic and pollen tube growth experiments; *M. guttatus* was a maternal recipient only in the transcriptomic study.

To evaluate PMPZ interactions outside the context of classical self-incompatibility systems, we restricted our focal taxa to self-compatible species, with the exception of *Castilleja* spp., which are partially self-incompatible (Hersch and Roy 2007). The condensed growing season and high local species richness at RMBL result in frequent interspecific pollen transfer, making this system well suited for examining pollen-pistil interactions across evolutionary distance.

### Semi-*in vivo* pollination assays and RNA sampling

To characterize pistillar transcriptional responses to pollen identity, we conducted *semi-in vivo* pollen tube growth assays following Qin et al. (2009) and (Cohen et al. 2025a). *P. groenlandica* and *M. guttatus* pistils were pollinated with eight and five pollen donors, respectively (**Fig. 1**), spanning closely and distantly related taxa. Fewer pollen types were used for *M. guttatus* due to limited availability of close relatives near RMBL. All pollen was collected on the day of use and stored at 4°C prior to pollination.

**Fig. 1.**
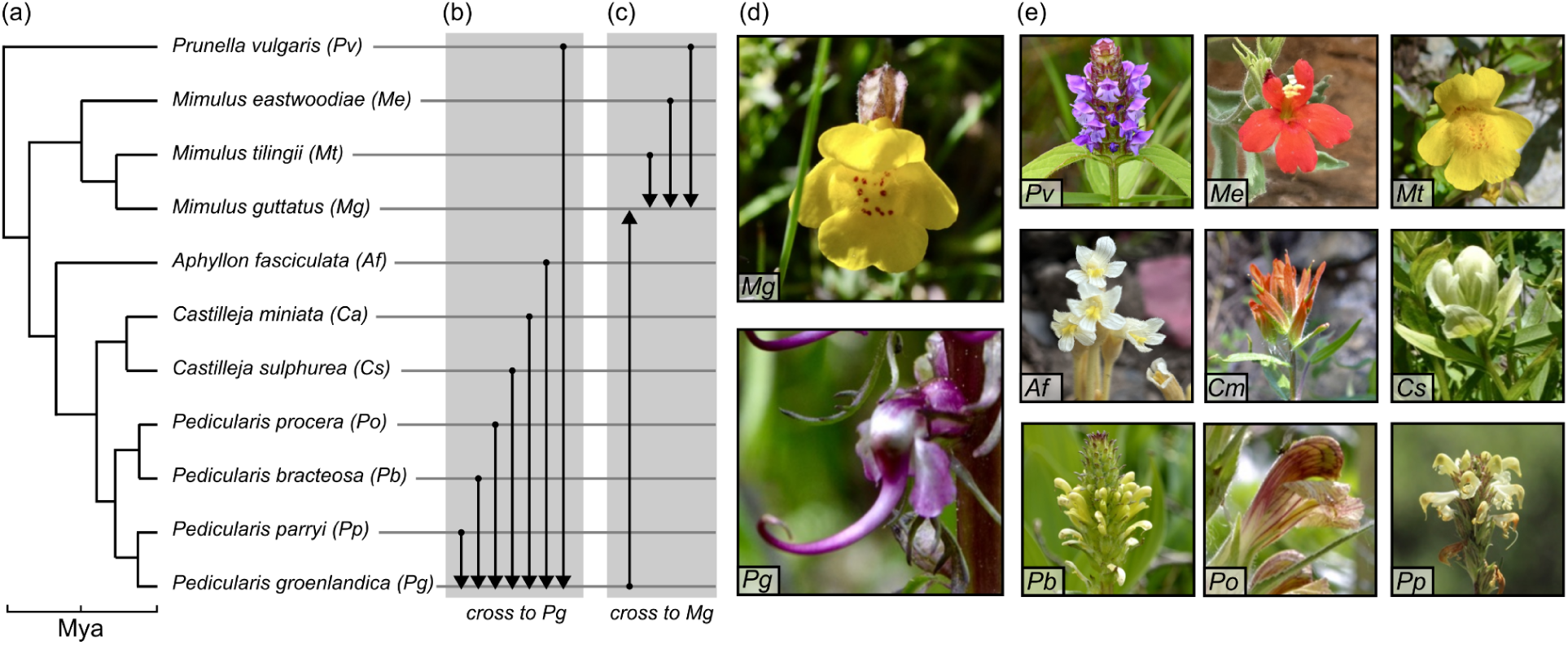
Species included in this study. a) Time-calibrated phylogeny based on ITS sequences of the species included, with time in millions of years (Mya) denoted on the x-axis. b) Species crossed to *P. groenlandica* in either the transcriptome or PTGR study denoted with arrows. c) Species crossed to *P. groenlandica* in either the transcriptome or PTGR study denoted with arrows. d) *M. guttatus* (*Mg*) and *P. groenlandica* (*Pg*) were used as maternal, pollen recipient species. e) Heterospecific pollen donor species.

Late-stage unopened buds (post-anther dehiscence, pre-anthesis) were collected to ensure virgin stigmas. We tested stigma receptivity using hydrogen peroxide on five buds in each collection batch. If all five were determined to be receptive, then the batch was deemed suitable for pollination assays. Pistils were excised, truncated to 2 mm, and stigmas verified to be pollen-free under 40× magnification. Each pistil then received a single-species pollen load. Pollinated pistils were placed on pollen tube growth medium (Cohen et al. 2025b), with the ovary positioned within 1 mm of the truncated pistil to promote continued tube growth across the cut interface (**Fig. 2a**). All pollen donors were tested for *in vitro* germination in the same medium prior to experimentation, and all exhibited >80% germination rates.

**Fig. 2.**
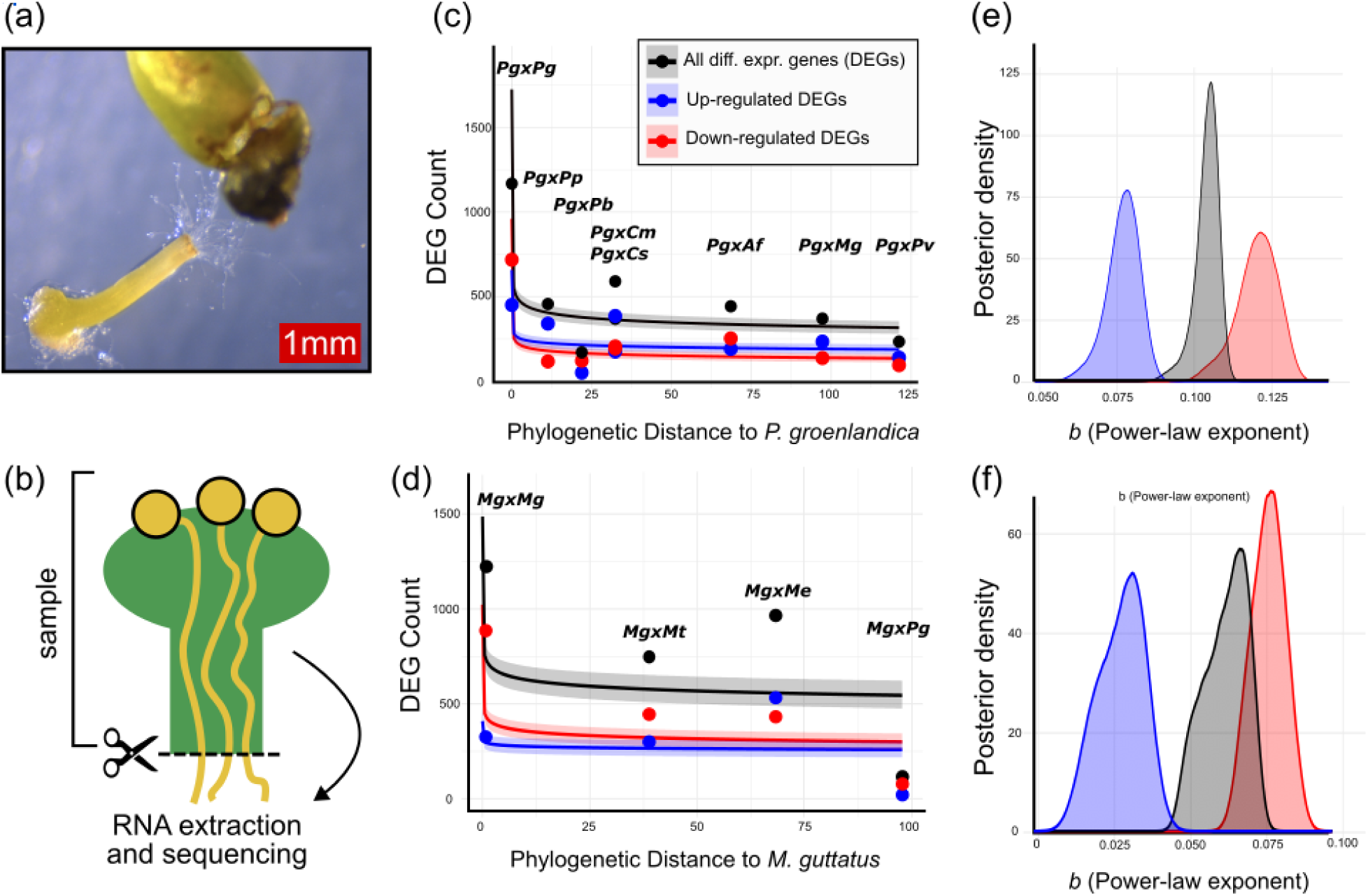
The number of genes differentially expressed in response to pollination decays with genetic distance. a) RNA was sampled from semi-*in vivo* pollen tube growth assays, conducted on petri dishes. b) We sampled only the pistil tissue for RNA sequencing, cutting off the protruding pollen tubes prior to sampling. c) Differentially expressed genes (DEGs) in crosses to *Pedicularis groenlandica*, including up-regulated (blue), down-regulated (red), and all DEGs (black). We fit a negative power-law model and show the predicted curves for up-regulated, down-regulated, and all DEGs are mapped over the data. Phylogenetic distance (horizontal axis) is patristic phylogenetic distance calculated using the time-calibrated phylogeny. d) Negative power law model fit to DEG data from crosses to *M. guttatus*, excluding the *M. guttatus* X *P. vulgaris* outlier. e) Posterior predictive curves for the b parameter (negative power-law exponent) in all models for crosses to *P. groenlandica*, shown separately for up-regulated, down-regulated, and all DEGs. In each case, the estimate for the b parameter is >0, indicating a significant negative relationship between phylogenetic distance and the number of DEGs. f) As in *e*, but for the model fit to crosses to *Mimulus guttatus*.

Assays were incubated under controlled growth chamber conditions (18 h light at 23°C, 6 h dark at 18°C). After 24h, pistils with visible pollen tubes emerging from the truncated end were collected to ensure that RNA sampling included active pollen tube growth within the style rather than pollen deposition on the stigma alone. Pollen tubes were removed from the cut end to minimize male tissue contribution (**Fig. 2b**), and pistils were flash-frozen in a dry ice-ethanol slurry and stored at −80°C. Because extractions were conducted under field conditions, liquid nitrogen was not available and all flash-freezing steps used dry ice-ethanol. Unpollinated controls underwent identical handling but received no pollen.

Three pistils were pooled per extraction, and three biological replicates were collected per pollination treatment. RNA was extracted using the Qiagen RNeasy Plant Mini Kit. Due to low input tissue, libraries were prepared using SMART-seq v4 ultra low-input protocols and sequenced on an Illumina MiSeq platform (Admera Health, NJ, USA).

### Differential expression quantification

Raw reads were trimmed for adapter sequences and low-quality bases (<Q30) using Trim Galore (Krueger 2015). Ribosomal RNA reads were removed with Ribodetector (Deng et al. 2022), and sequence quality was assessed using FastQC (Andrews 2010). Filtered reads were aligned to the respective reference genomes for *P. groenlandica* (Cohen et al. 2025a) and *M. guttatus* (NONTOL v4.0, DOE-JGI) using STAR v2.7.11 (Dobin et al. 2013), and gene-level read counts were generated with featureCounts v2.0.6 (Liao et al. 2014).

Differential expression analyses were performed separately for *P. groenlandica* and *M. guttatus* using DESeq2 v1.42.1 (Love et al. 2014). For each maternal species, expression in each pollination treatment was compared to its corresponding no-pollen control. Model fit and replicate clustering were evaluated using principal component analysis and sample-to-sample distance heatmaps. One *P. bracteosa × P. groenlandica* replicate exhibited aberrant clustering and was removed prior to downstream analyses. Genes were considered differentially expressed if they showed an absolute log₂fold-change > 2 and an adjusted p-value < 0.05. Differentially expressed gene (DEG) sets from each cross were used for subsequent comparative and modeling analyses.

### Modeling the effect of phylogenetic distance on pollen–pistil interactions

To quantify evolutionary distance among study species, we reconstructed a phylogeny using ITS sequences downloaded from NCBI (**Table S1**). Sequences were aligned using the R package *msa* (Bodenhofer et al. 2015), and a maximum likelihood tree was inferred in *phangorn* (Schliep 2011), with optimization starting from an initial neighbor-joining tree. The resulting topology was consistent with published phylogenetic relationships for these clades. The tree was transformed into an ultrametric, time-calibrated phylogeny by fitting a relaxed clock model under penalized likelihood using the *chronos* method in *ape* (Paradis and Schliep 2019), with internal node age calibrations for the divergence times between plant families from TimeTree.org (Kumar et al. 2022). We performed 1,000 independent calibration iterations and averaged branch lengths to generate the final tree. Patristic distances between each pollen donor and the maternal recipients (*P. groenlandica* and *M. guttatus*) were calculated using *adephylo* (Jombart et al. 2010). These distances were used as predictors in downstream models.

### Modeling phylogenetic effects on differential expression

We then evaluated how phylogenetic distance predicts transcriptional response using two complementary modeling approaches. First, we modeled the number of DEGs observed in each pollination treatment as a function of phylogenetic distance to the maternal species. Because the number of pollination treatments per maternal species was limited (<10), our objective was to compare plausible functional relationships rather than infer a precise evolutionary decay model. We compared negative Hill, exponential, and power-law functions under both Poisson and negative binomial error structures within a Bayesian framework, controlling for library size across samples. Model performance was evaluated using RMSE, WAIC, BIC, and LOOIC (**Table S2**). Across criteria, a Poisson-distributed negative power-law function provided the most consistent fit and was subsequently applied to both maternal species datasets.

Second, we modeled gene-level expression patterns across pollination treatments to *P. groenlandica* within a hierarchical Bayesian framework analogous to joint species distribution models. Variance-stabilized counts were obtained using the DESeq2 *vst* function (Love et al. 2014), and expression for each gene was modeled as a function of phylogenetic distance to the maternal species while controlling for RNA-seq library size. To evaluate whether gene-specific properties modulated distance effects, we incorporated gene length, GC content, and co-expression network membership as predictors. Co-expression networks were inferred using WGCNA v1.73 (Langfelder and Horvath 2008) based on TPM values across samples, following (Gilman et al. 2022) and (Cohen et al. 2025a). Models were implemented using the R package *Hmsc* (Tikhonov et al. 2020).

### Differentially expressed gene identity and function

To characterize the functional composition of transcriptional responses, we assigned Gene Ontology (GO) annotations to differentially expressed genes (DEGs). For each maternal species, protein sequences were matched to the closest *Arabidopsis thaliana* orthologs (Cheng et al. 2017) using BLASTp (Ye et al. 2006). Enriched GO Biological Process terms were identified for each pollination treatment using the *enrichGO* function in *clusterProfiler* (Wu et al. 2021); GO terms were classified as enriched if p<0.05 and q<0.05. We compared GO enrichment profiles among conspecific and heterospecific crosses spanning increasing phylogenetic distance.

To examine how DEG composition varied across divergence, we quantified overlap in DEG sets across nested clades of pollen donors. We identified genes unique to conspecific crosses, genes shared among intrageneric crosses, and genes retained across progressively broader phylogenetic groupings, allowing us to determine whether particular gene subsets were conserved or lost as divergence increased.

To assess whether pollination responses involved the same genes between the two maternal species used in our study, we inferred orthogroups among *P. groenlandica*, *M. guttatus*, and *A. thaliana* using OrthoFinder (Emms and Kelly 2019). We evaluated whether DEGs identified in analogous cross types (conspecific, intrageneric, intergeneric, and interfamily crosses) belonged to the same hierarchical orthogroups (HOGs). We tested whether DEGs in our dataset overlapped with *A. thaliana* genes experimentally validated to function in “pollination”, “pollen-pistil interactions”, “pollen tube growth”, or “pollen germination”. Gene lists were obtained from the TAIR database and mapped to orthogroups to identify shared functional categories.

### Pollen tube growth rate quantification

To quantify pollen tube growth rates (PTGRs), we conducted experimental field pollinations using *P. groenlandica* as the primary maternal species. We pollinated *P. groenlandica* with conspecific pollen and with pollen from five heterospecific donors (*P. bracteosa*, *P. procera*, *Castilleja miniata*, *C. sulphurea*, and *M. guttatus*). In parallel, we measured conspecific PTGRs within each donor species to enable direct comparison between heterospecific growth in *P. groenlandica* pistils and conspecific growth in the donor’s native pistil. Although *P. parryi* was included in transcriptomic analyses, it was excluded from the PTGR experiment as it is a high altitude species (growing only at altitudes above 11,500ft; (Macior 1995), making it less accessible for field-based crosses.

Measuring PTGRs requires sampling pollen tubes growing within pistils at two timepoints following pollination. To determine appropriate sampling timepoints, we established the time required for conspecific pollen grains to germinate and pollen tubes to traverse the stigma using both field-based and *semi-in* vivo assays. For field-based measurements, we pollinated five individuals in the morning, then returned for collection after 4, 6, 8, 10, and 12 hours. Pistils were fixed, cleared, stained with fuchsin and acetocarmine, and examined microscopically to determine when pollen tubes approached the base of the stigma. For semi-*in vivo* crosses, we set up five conspecific cross assays per species and observed the assays under a dissection microscope at 40X magnification every hour, recording whether pollen tubes could be observed growing through the truncated end of the pistil. Based on these preliminary measurements, we selected two collection timepoints per species that captured active pollen tube growth within the style prior to tube arrival at the base (**Table S3**).

For the main experiment, individuals were bagged prior to anthesis. We returned to these individuals approximately two weeks later to experimentally pollinate the flowers. We collected pollen from ∼30 unbagged individuals each morning prior to hand pollination, removed the anthers, and vortexed the anthers to create a homogenous mixture of pollen for *P. groenlandica* and the five heterospecific treatment species. All conspecific crosses, therefore, were completed with a mixture of outcross pollen from distinct individuals.

Pollinations were conducted in the morning under consistent environmental conditions (33-35°F, sunny), and conspecific and heterospecific pollinations were performed on the same individuals and on the same day for each donor type to minimize environmental effects (Taylor and Hepler 1997). In *P. groenlandica*, four flowers per individual were pollinated – two with conspecific pollen and two with heterospecific pollen from a single donor species. Pistils were collected at approximately 8 h (T1) and 9.5 h (T2) post-pollination, with exact times of pollination and collection recorded. Thirty maternal individuals were sampled per heterospecific treatment, and unpollinated controls were sampled. Conspecific pollinations in donor species were conducted concurrently, and we additionally collected data from 30 individuals. Pistils which did not receive experimental pollen treatments were additionally collected as a control comparison from each individual.

Fixed pistils were processed following Cohen et al. (2025b). Samples were washed, softened in NaOH, stained with 0.1% aniline blue in 0.1 M K₃PO₄, and imaged using a Zeiss LSM700 epifluorescence microscope (350 nm excitation, 450 nm emission; **Fig. S1**). Observers were blind to treatment and timepoint during imaging. For each pistil, the longest pollen tube was measured from the stigma surface to its distal tip using ZENLite software. Style length and stigma depth were also recorded, as both may influence tube growth. Samples in which pollen tubes had reached the base of the pistil were excluded because growth time could not be reliably determined.

Microscope images with poor visibility (showing blurry, broken, tangled, or cut-off styles) were removed from the dataset (n=120 pistils). We additionally removed individuals in which we observed pollen tubes in the no pollen controls (124 pistils, 32 individuals) and samples where pollen tubes grew to the end of the pistil (n=66 individuals). From the remaining samples (n=519 pistils), we assembled 109 intact paired T1-T2 measurements. We initially calculated strict PTGRs using paired measurements (Δdistance/Δtime; **Fig. S2**), but restricting analyses to paired samples substantially reduced statistical power. We therefore calculated PTGR as pollen tube length divided by time since pollination, treating T1 and T2 samples independently. This metric captures both time to germination and pollen tube growth timing, and therefore still reflects overall pollen performance within the pistil; hereafter, we refer to this measure as PTGR.

### Statistical analysis of pollen tube growth

To test whether phylogenetic distance predicts PTGR in *P. groenlandica* pistils, we fit a Gamma-distributed generalized linear model with a log link function, which is appropriate for strictly positive rate data. Phylogenetic distance between pollen donor and maternal species was included as a predictor, and style length and stigma depth were included as covariates to account for morphological variation among pistils. We additionally conducted pairwise comparison of each treatment group (conspecific, intrageneric, intrafamily, and interfamily crosses) using a Chi-Square test with Tukey correction. We further compared heterospecific PTGRs to conspecific PTGRs in donor species using pairwise two-tailed Welch’s t-tests. Together, these analyses allowed us to evaluate both the effect of evolutionary distance on pollen performance within a common maternal background and the relative performance of pollen across maternal environments.

## Results

### Pollination response by phylogenetic distance

After quality trimming and rRNA removal, we obtained an average of 24.9 million reads per replicate per cross in *Pedicularis* libraries, of which a mean of 16.0 million reads (64.7%) uniquely mapped to the *P. groenlandica* genome. This rate was relatively low because pistils contained both maternal and paternal (pollen tube) tissue and heterospecific reads are expected to map to the genomes at a lower rate. As anticipated, the highest mapping rate occurred in the conspecific *P. groenlandica × P. groenlandica* cross (86.0%). In *M. guttatus*, we obtained an average of 23.4 million reads per replicate per cross, with a mean of 12.6 million reads (53.8%) uniquely mapping. Similarly, the highest mapping rate occurred in the conspecific *M. guttatus × M. guttatus* cross (75.3%).

Across all crosses to *P. groenlandica*, we identified 3,796 differentially expressed genes (DEGs), whereas 6,199 DEGs were identified among crosses to *M. guttatus*. Much of this difference was driven by an exceptionally large response in the *M. guttatus × P. vulgaris* cross, which alone accounted for 3,093 DEGs. In all other crosses, the number of DEGs was comparable between maternal species.

In crosses to *P. groenlandica*, the number of DEGs declined with increasing phylogenetic distance according to a negative power-law relationship (**Fig. 2c,e**). This pattern held when considering all DEGs combined and when analyzing up- and down-regulated genes separately. In contrast, crosses to *M. guttatus* initially showed little relationship between DEG count and phylogenetic distance due to the large number of DEGs in the most distant cross (*M. guttatus × P. vulgaris*; **Fig. S3**). Most DEGs in this cross were down-regulated, resulting in a weak relationship when considering only up-regulated genes. However, when the *P. vulgaris* cross was excluded, the relationship between DEG count and phylogenetic distance closely mirrored that observed in *P. groenlandica* (**Fig. 2d,f**).

At the gene level, our hierarchical model revealed an overall negative relationship between phylogenetic distance and differential expression in crosses to *P. groenlandica*. The beta coefficient associated with phylogenetic distance was negative for 78.4% of genes (**Fig. S4**), indicating that most genes were less likely to be differentially expressed as evolutionary distance increased. The negative relationship between phylogenetic distance and gene-level expression was not associated with gene length or GC content, but varied significantly with several co-expression modules (**Table S4**). Of the modules with non-zero gamma values modulating the relationship between phylogenetic distance and differential expression in our hierarchical model, most exhibited positive effect sizes, indicating that genes within those modules were more likely to be differentially expressed in more distantly related crosses. These modules were primarily associated with general cellular functions such as photosynthesis, plastid function, metabolism, and chromatin organization. In contrast, two modules with negative effect sizes were enriched for genes involved in hormone and stress signaling, indicating that signaling-related genes were less likely to be differentially expressed as genetic distance increased.

### Pollen tube growth rates across evolutionary distance

Pollen tube growth rates (PTGRs) declined significantly with increasing phylogenetic distance in *P. groenlandica* pistils (slope = −4.831 × 10⁻³, P = 0.022; **Fig. 3a**). Heterospecific PTGRs were slower than conspecific *P. groenlandica × P. groenlandica* PTGRs and were also slower than conspecific PTGRs measured within the donor species (**Fig. 3b**). Notably, the slowest growth rates occurred in intrageneric crosses (*P. groenlandica × P. bracteosa* and *P. groenlandica × P. procera*; **Fig. 3b**), rather than in the most phylogenetically distant crosses.

**Fig. 3.**
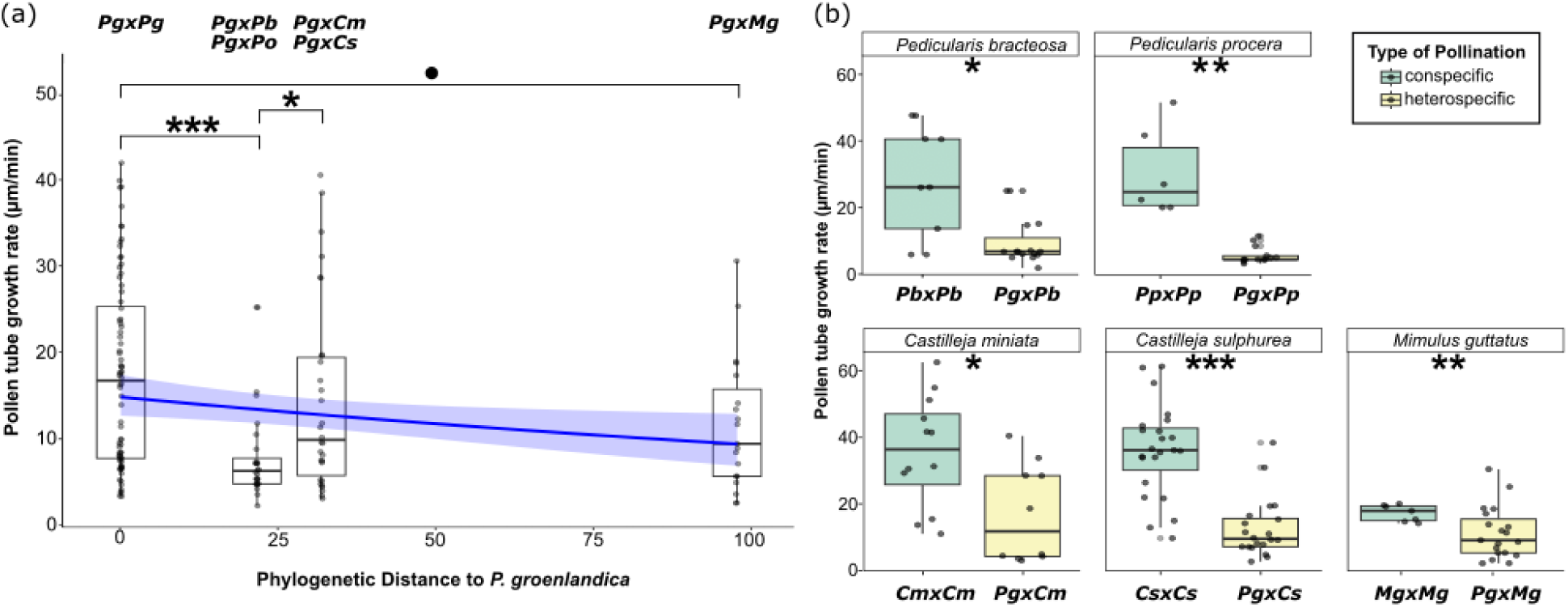
Pollen tube growth rates are slower in heterospecific crosses compared to conspecific crosses. a) In crosses to *Pedicularis groenlandica*, pollen tube growth rate decays with phylogenetic distance according to a gamma-distributed generalized linear model (slope=-4.83e^−3^±2.081e^−3^, *P*=0.02). Phylogenetic distance was calculated as patristic phylogenetic distance to *P. groenlandica* using the time-calibrated phylogeny. b) Comparison of heterospecific PTGRs in *P. groenlandica* (yellow) to conspecific PTGRs in each of the pollen donor species (green). Statistical differences among the cross categories is indicated above the boxplots as • (P<0.1), * (P<0.05), ** (P<0.01), and *** (P<0.001).

### Composition and functional identity of differentially expressed genes

Although the magnitude of transcriptional response was broadly similar between *P. groenlandica* and *M. guttatus* (excluding the *P. vulgaris* outlier), the identities and functional categories of DEGs differed substantially between maternal species and across phylogenetic distance. In crosses to *M. guttatus*, enriched GO biological process terms in the conspecific cross closely resembled those in intrageneric crosses (*M. tilingii* and *M. eastwoodiae*; **Fig. 4b**). These terms included processes directly related to pollen-pistil interactions, such as “unidimensional cell growth,” “cell wall biogenesis,” “pollination,” and “pollen tube development.” In more distant crosses (*M. guttatus × P. groenlandica* and *M. guttatus × P. vulgaris*), although some pollination-related genes remained differentially expressed, enriched terms increasingly reflected general cellular processes such as “regulation of transcription by RNA polymerase II,” “cellular response to organic cyclic compound,” and “response to nitrogen compound.” The shift from pollination-specific to background cellular processes across evolutionary divergence occurred even when the *P. vulgaris* cross was included, in contrast to the DEG count analysis, where that cross obscured distance-dependent patterns.

**Fig. 4.**
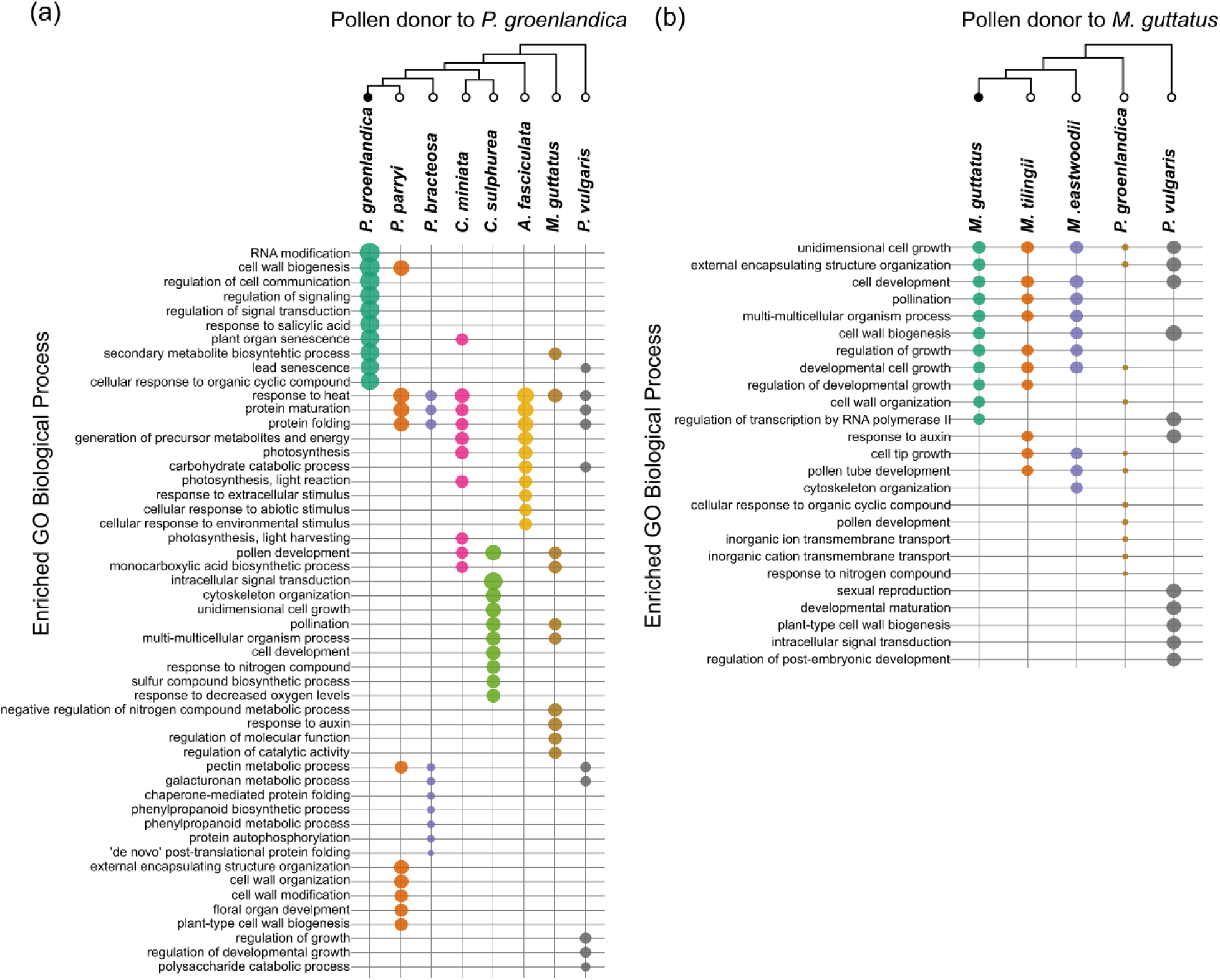
Variation in the identities of the DEGs in each cross. a) Top 10 GO terms enriched in the DEGs of each cross to *Pedicularis groenlandica* and b) to *Mimulus guttatus* (right). The relationship of the pollen donor species to *P. groenlandica* and *M. guttatus* is shown in the phylogeny above each plot, with the maternal species denoted with a black circle at their terminal nodes. The count of genes associated with each biological process term is indicated by the size of the dots.

Crosses to *P. groenlandica* exhibited a different pattern. Across all crosses, DEGs included both pollination-related and non-pollination-related processes (**Fig. 4a**). Intrageneric crosses (*P. parryi* and *P. bracteosa*) did not strongly resemble the conspecific cross in their GO profiles, though they were relatively similar to each other. Considerable variation was observed in the top enriched GO terms even among closely related donor species (e.g., *C. miniata* and *C. sulphurea*). Notably, “response to heat” was among the top enriched GO terms in six of seven heterospecific crosses to *P. groenlandica*, and “protein maturation” and “protein folding” were enriched in five of seven crosses. Some GO terms were shared between crosses to *P. groenlandica* and crosses to *M. guttatus*, particularly those related to cell wall processes (“cell wall biogenesis,” “plant-type cell wall biogenesis,” “cell wall organization”) and pollen-related processes (“pollination,” “pollen development,” “unidimensional cell growth”).

As phylogenetic breadth increased in DEG overlap analyses, the number of shared genes declined (**Fig. 5a-b**). A large proportion of DEGs were therefore unique to individual heterospecific crosses, particularly at greater evolutionary distances. Although this pattern was partially driven by the overall decay in DEG number with distance, it persisted in crosses to *M. guttatus* even when the *P. vulgaris* cross—despite containing over 3,000 DEGs—was included (**Fig. 5b**).

**Fig. 5.**
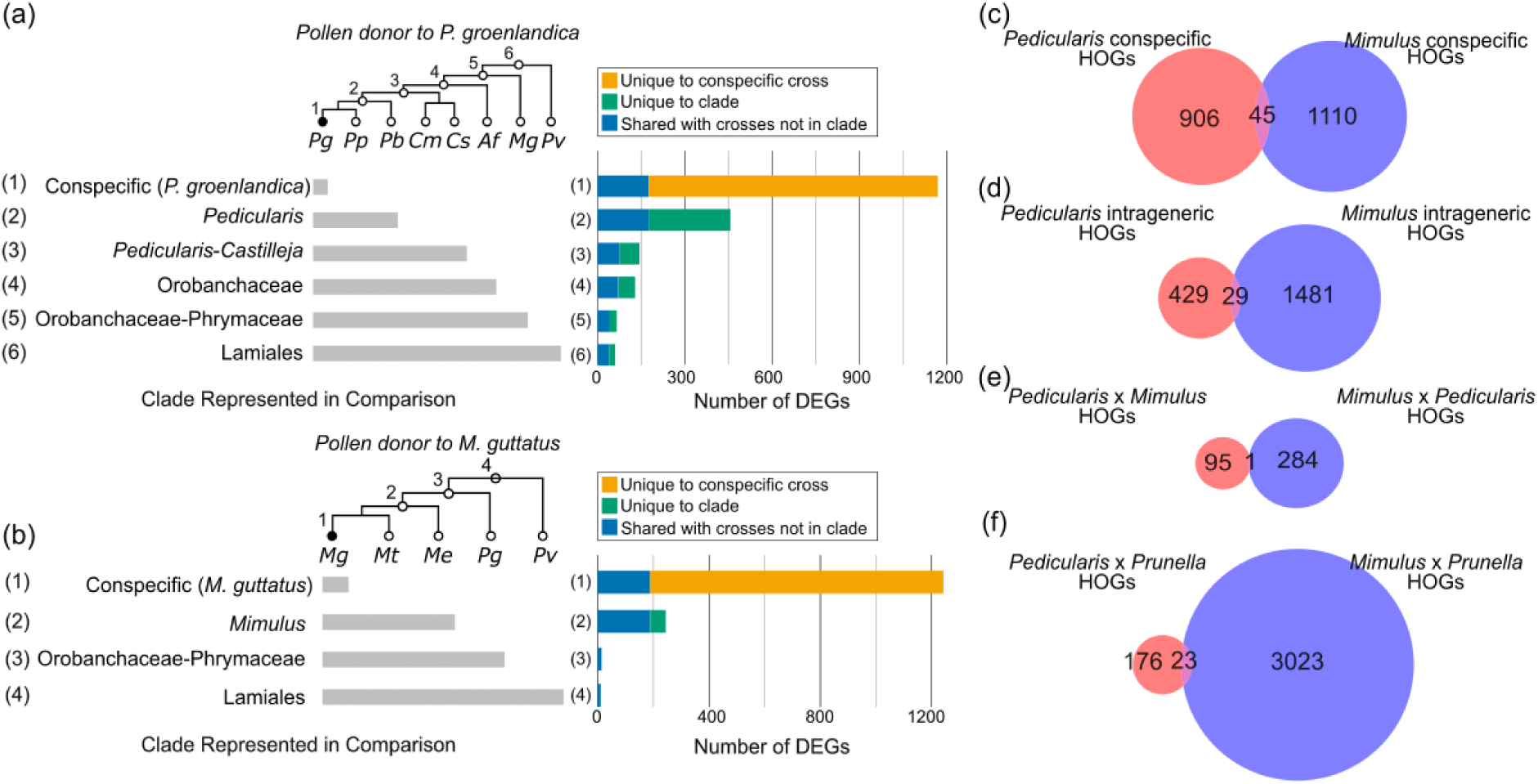
Lineage specificity in pollination responses. a-b) Comparison of the DEGs in increasingly large clades across the phylogenetic breadth of species included in this study. The species included in each successive comparison are denoted by the phylogeny and grey bars to the left of the plot, where the grey bars overlap with the species included in each clade. In each plot, the conspecific cross is represented by the topmost bar, and the DEGs unique to the conspecific cross (orange) are compared to the DEGs shared with any heterospecific cross (blue). The bars for heterospecific crosses represent increasingly large clades, and show the DEGs that are uniquely expressed in that clade (green) compared to DEGs shared with either the conspecific cross or with other heterospecific crosses not included in that clade (blue). We show that the number of DEGs shared across all members of the clade decreases as the clade grows to include crosses to more distant relatives in crosses to both a) *P. groenlandica* and b) *M. guttatus*. c-f) Venn diagrams showing overlap orthology among DEGs in crosses to *P. groenlandica* and *M. guttatus*, comparing analogous cross-types to each maternal species.

### Lineage specificity of pollination responses

To evaluate conservation of pollination responses across maternal species, we compared orthology among DEGs using hierarchical orthogroups (HOGs). Surprisingly, even conspecific pollination responses were highly lineage-specific (**Fig. 5c**). Only a single shared HOG was identified between reciprocal intergeneric crosses (*P. groenlandica × M. guttatus* and *M. guttatus × P. groenlandica*), representing the lowest overlap among comparisons (**Fig. 5e**). Similarly, despite the large number of DEGs in the *M. guttatus × P. vulgaris* cross, only 23 shared HOGs were identified relative to other crosses (**Fig. 5f**), a magnitude comparable to other pairwise comparisons.

To further assess lineage specificity, we examined overlap between our DEGs and experimentally validated pollination-related genes in *Arabidopsis thaliana* (**Table S5**). Of 5,146 annotated *A. thaliana* pollination genes, only 754 had orthologs in *P. groenlandica* or *M. guttatus* that were differentially expressed in any cross. On average, 17.7% (SD = 0.08) of DEGs were orthologous to *A. thaliana* pollination genes. The proportion of DEGs overlapping *A. thaliana* pollination genes did not decline with genetic distance in either maternal species. Notably, the *M. guttatus × P. vulgaris* cross exhibited the fewest *A. thaliana* pollination orthologs despite having the largest total number of DEGs.

## Discussion

### Molecular responses to pollination weaken with evolutionary divergence

In *P. groenlandica*, phylogenetic distance negatively predicts the molecular response (**Fig. 2c–f**). This pattern is notable because heterospecific pollen tubes can grow through pistils and achieve fertilization across wide phylogenetic distances in the same alpine community (Cohen et al. 2025b). Together, these observations suggest that distantly related heterospecific pollen often fails to elicit a strong, pollination-specific pistillar response, even when pollen tube growth is physiologically possible. Rather than activating a generalized rejection mechanism against increasingly distant heterospecific donors, coordinated support of pollen tube growth by the pistil appears to decline as evolutionary divergence increases. This interpretation contrasts with the hypothesis that pistil-acting PMPZ mechanisms should broadly arrest heterospecific pollen tube growth as a defense against interspecific pollen transfer (Ashman and Arceo-Gómez 2013; Hamlin et al. 2017; Streher et al. 2020), for which empirical support remains mixed (Arceo-Gómez et al. 2019; Moreira-Hernández and Muchhala 2019; Cohen et al. 2025b).

A broadly similar decay pattern was evident in crosses to *M. guttatus* once the *M. guttatus × P. vulgaris* cross was excluded. That cross produced an unusually large transcriptional response (3,093 DEGs), flattening the distance relationship when all treatments were considered together. We evaluated several non-biological explanations for this outlier, including library size and pollination identity. Library sizes were comparable to other treatments, and mapping reads to the *P. vulgaris* genome suggested that the paternal component was consistent with the intended donor, at rates comparable to other heterospecific crosses and to the *P. groenlandica × P. vulgaris* cross. We also do not expect that *M. guttatus* experiences unusually frequent exposure to *P. vulgaris* pollen relative to other donors in our sampling design, which makes an adaptive explanation less compelling. As a result, we cannot currently identify a confident explanation for the exceptional DEG count in the *P. groenlandica × P. vulgaris* cross. There is some evidence that *Mimulus* species can exhibit stronger transcriptional responses to heterospecific than conspecific pollination, although existing work has focused on congeneric comparisons (M. Mattson, *personal communication*). Because we did not measure *P. vulgaris* pollen germination or tube growth in *M. guttatus* pistils, we also cannot yet evaluate whether this cross represents an unusual physiological interaction relative to the other donors used here.

Although total DEG counts were the most direct signal of distance dependence in *P. groenlandica*, the functional composition of these genes provided an additional axis of change. In crosses to *M. guttatus*, pollination-associated GO categories were most prominent in conspecific and intrageneric crosses and declined with increasing phylogenetic distance, particularly in the *M. guttatus × P. groenlandica* cross. This pattern complements the DEG-count decay (excluding the *P. vulgaris* outlier) and suggests that the “pollination-specific” component of the response attenuates with divergence even when a transcriptomic response remains detectable. In crosses to *P. groenlandica*, GO enrichment was more variable across donors, and a single monotonic functional trend was less apparent. One repeated signal stood out: “response to heat” was enriched in nearly all heterospecific crosses. In *A. thaliana*, incompatible pollen can elicit a general stress response (Kodera et al. 2021), and heat shock pathways can be induced by diverse stressors beyond temperature (Swindell et al. 2007). More broadly, ROS and calcium signaling are central to both stress physiology and pollen-pistil interactions (Steinhorst and Kudla 2013; Duan et al. 2014; Kaya et al. 2014; Devireddy et al. 2021), and ROS was recently implicated in a lineage-specific mechanism that suspends heterospecific pollen germination in Brassicaceae (Cao et al. 2025). Notably, Cao et al. (2025) found no clear orthologs of their key genes across a broad angiosperm panel, suggesting that while stress-related signaling may recur as a physiological theme, the specific genes involved may differ among lineages.

Our transcriptomic results suggest that pistillar response to pollination is strongest for conspecific pollen and attenuates with evolutionary divergence. Pollination-relevant functional categories were most prominent in conspecific and intrageneric crosses. Although some background cellular and stress-associated genes exhibited increased expression with increasing phylogenetic distance, the expression of a majority of genes was inversely related to phylogenetic distance. Furthermore, genes associated with functional categories associated with signaling, a major component of successful pollen-pistil coordination, were less commonly represented in crosses among distant relatives. These patterns are more consistent with decay of coordinated pistil support than with the existence of a broadly conserved, distance-escalating heterospecific rejection program. In this framework, distantly related pollen may elicit weaker and less specialized transcriptional responses, even when tube growth remains physiologically possible, as demonstrated previously in this system (Cohen et al. 2025b).

### Heterospecific pollen tubes grow more slowly, but close relatives are a special case

Consistent with reduced maternal support of heterospecific pollen, pollen tube growth rates (PTGRs) in *P. groenlandica* pistils declined with increasing phylogenetic distance (**Fig. 3a**). Moreover, heterospecific PTGRs were consistently slower than conspecific PTGRs measured both in *P. groenlandica* and in the donor species’ native pistils (**Fig. 3**). The concordance between declining pollination-associated transcriptional signatures and reduced PTGR raises the possibility that diminished molecular coordination could translate into measurable physiological consequences.

However, the slowest PTGRs did not occur in the most distantly related crosses. Instead, the most pronounced reductions were observed in intrageneric crosses (*P. groenlandica × P. bracteosa* and *P. groenlandica × P. procera*). Closely related heterospecific donors also tended to elicit relatively larger transcriptomic responses than distant donors, both in DEG number and in the prevalence of pollination-associated functional categories. Importantly, the overall regression of PTGR on phylogenetic distance remained negative and statistically significant despite the intrageneric minimum, indicating that the slowdown among close relatives represents an additional deviation layered onto a broader distance-dependent decline. Together, slower tube growth paired with an elevated gene expression response in close relatives suggests that, in addition to reduced support for heterospecific pollen in general, a distinct mechanism may operate to slow or impede pollen tube growth specifically in within-genus heterospecific crosses. This does not contradict the broader trend of reduced recognition and support with divergence, but instead suggests that close relatives may represent a special case in which maternal tissues both fail to provide full conspecific-like support and also activate responses that further hinder tube performance.

The existence of a mechanism that strongly affects close relatives is consistent with the expectation that heterospecific pollen transfer among close relatives can impose the strongest fitness costs because it is most likely to result in hybridization or to create intense competition within the transmitting tract. This logic likely contributes to why many studies on reproductive interference and heterospecific pollen transfer have focused on congeners (Eaton et al. 2012; Baek et al. 2015; Arceo-Gómez et al. 2019; Moreira-Hernández and Muchhala 2019; Rifkin et al. 2023). In our self-compatible study set, the intrageneric growth-rate minimum suggests that, independent of classical self-incompatibility, selection imposed by closely related heterospecific pollen may favor mechanisms that reduce heterospecific tube performance. We use “reduce” rather than “suspend” because pollen tube passage through pistils is possible even across deep divergence in this system (Cohen et al. 2025b). Although our data do not identify causal pathways, the repeated enrichment of stress-related GO terms such as “response to heat,” together with known roles for ROS and calcium signaling in both pollination and stress responses, makes these pathways plausible candidates for closer mechanistic work.

### Pollination-response genes are strongly lineage-specific

Across maternal species, the genes involved in pollination responses were surprisingly lineage-specific. The majority of DEGs in crosses to *P. groenlandica* were not orthologous to DEGs in crosses to *M. guttatus*, and only a small fraction of DEGs in either lineage were orthologous to well-characterized *A. thaliana* pollination genes. This was unexpected, particularly for conspecific crosses, in which we expected to observe differential expression of genes that are broadly characterized as being associated with a pollination response. Much of the mechanistic framework linking functional annotations for pollen germination, pollen tube growth, and pollen-pistil signaling has been developed from a small set of model taxa (predominantly *A. thaliana*, *Zea mays*, *Nicotiana tabacum*, and *Solanum lycopersicum* (Herrero and Hormaza 1996; Figueroa-Castro and Holtsford 2009; Qin et al. 2009; Liu et al. 2012; Duan et al. 2014; Baek et al. 2015; Alves et al. 2019). One interpretation of our results is that conspecific pollination responses may involve a conserved core set of processes while recruiting many lineage-specific genes and regulators, such that the transcriptomic signature of “pollination response” differs substantially among angiosperm lineages. Because we only sampled conspecific pollination in two maternal species, broader phylogenetic sampling of conspecific responses will be required to test how general this pattern is. Within the scope of this study, the most direct implication is that differences between conspecific and heterospecific responses may often be mediated through lineage-specific networks rather than through a universal set of conserved “pollination genes.”

Whereas we expected at least partial conservation in genes responding to conspecific pollination, we did not expect a similarly conserved transcriptomic program for heterospecific crosses. In systems where self-incompatibility (SI) contributes to interspecific barriers, the molecular components underlying pollen rejection are relatively well-characterized and conserved (Broz and Bedinger 2021), offering a model of how heterospecific rejection can be genetically specified. Here, however, we focused on self-compatible taxa to explicitly examine PMPZ interactions outside classical SI. In this framework, one prediction for the relationship between pistillar transcriptomic response and evolutionary distance comes from the incongruence hypothesis (Hogenboom et al. 1997), which posits that incompatibilities arise as interacting pollen-pistil components diverge via neutral mutations, producing lineage- and pairing-specific mismatches. Importantly, this framework predicts that a breakdown of coordination should occur gradually as divergence accumulates, rather than as a simple binary switch between “compatible” and “incompatible” states. Consistent with incongruence theory, we find that the magnitude of the stylar transcriptomic response decays quantitatively with evolutionary distance, and that DEG identity and functional composition vary continuously across donor lineages. Our results therefore provide empirical support for a graded, divergence-dependent model of pollen-pistil incongruence rather than a conserved heterospecific rejection program.

Recent work provides a useful example of how lineage specificity can arise even when a mechanism is clearly identified. Cao et al. (2025) described a PMPZ mechanism in self-incompatible Brassicaceae in which heterospecific pollen triggers ROS accumulation through SRK-interacting Interspecific Pollen Signal (SIPS), suspending pollen germination. They further reported limited homology of key components outside Brassicaceae, suggesting that the molecular basis of this barrier is restricted to the family. In our study, we likewise observe that heterospecific pollination responses vary both with maternal and paternal species, including substantial donor-to-donor variation in enriched biological processes for *P. groenlandica* crosses and increasing functional divergence among more distant crosses in *M. guttatus*. In other words, the molecular response depends on the specific pairing of pollen and pistil, consistent with lineage- and interaction-specific incongruence.

### Implications for PMPZ barriers and the fitness costs of heterospecific pollen

Our results suggest that PMPZ barriers in angiosperms can arise primarily through the decay of pistil-acting mechanisms that evolved to promote conspecific pollen tube growth, rather than through targeted recognition and arrest of heterospecific pollen. Across a broad phylogenetic scale, heterospecific pollinations elicited weaker transcriptomic responses and reduced PTGRs relative to conspecific crosses, yet pollen tubes can still reach the ovules under pure-load conditions (Cohen et al. 2025b). Combined, these observations are consistent with heterospecific tube growth being shaped by pollen-intrinsic behavior operating in a pistil environment that provides baseline permissiveness but limited active support. Indeed, Sanders and Lord (1989) showed that pistils can translocate inert particles such as latex beads, consistent with the idea that some aspects of downward movement through transmitting tissue may reflect general properties of the pistil environment, rather than explicit coordination between congruent pollen and pistils (Sanders and Lord 1989, 1992).

Beyond a general breakdown in heterospecific pollen–pistil compatibility, incongruence may also arise through the rapid divergent evolution of coevolving pollen and pistil genes. Processes such as pollen competition could impose strong positive selection on conspecific pollen–pistil interaction genes, driving rapid divergence in pollen recognition systems even among closely related taxa. This possibility is supported by our observation that many conspecific pollen–pistil interaction genes exhibit strong lineage specificity. Under this scenario, incongruence in heterospecific pollen–pistil interactions may ultimately reflect divergence in the molecular programs underlying conspecific pollen–pistil recognition.

A key implication is that the ability of heterospecific pollen tubes to grow does not imply they can compete effectively with conspecific pollen. This distinction matters for understanding fitness costs of heterospecific pollen receipt (Ashman and Arceo-Gómez 2013; Moreira-Hernández and Muchhala 2019). Under pure-load pollinations, we have evidence that heterospecific pollen tubes can reach the end of the pistil and achieve fertilization across wide divergence (Cohen et al. 2025b), while also showing that heterospecific tubes consistently grow more slowly than conspecific tubes (**Fig. 3**). In nature, however, pollen receipt often occurs as mixed loads, and seed set frequently declines as the heterospecific:conspecific ratio increases (Ashman and Arceo-Gómez 2013), including in alpine systems (Brosi and Briggs 2013; Briggs et al. 2016). Such mixed pollen loads may also elicit responses that differ fundamentally from those observed under pure-load conditions. Even a small amount of conspecific pollen could be sufficient to activate pistil support pathways that enhance pollen tube growth more generally, potentially extending that support to heterospecific tubes growing in the same transmitting tract. In agricultural systems, “mentor” or “pioneer” pollen has been used to facilitate fertilization by otherwise incompatible pollen types when applied sequentially (Visser 1981, 1983), demonstrating that pollen-pistil interactions can be modified by the order and identity of prior pollen deposition. If analogous dynamics occur under mixed heterospecific pollinations, ovule usurpation may become more likely than predicted from pure-load experiments, as heterospecific tubes could benefit from conspecific-triggered support. Testing this hypothesis will require the ability to track pollen donor identity while pollen tubes grow within the same pistil. Because aniline blue staining cannot distinguish pollen tubes by species, future work could employ differently colored variants of green fluorescent protein (GFP) to label donor pollen (Liu et al. 2012) or RNA-based genotyping approaches to assign pollen tubes to species of origin (Lobaton et al. 2021). Such approaches would allow direct quantification of donor-specific pollen tube growth dynamics under ecologically realistic mixed-load conditions.

## Data Availability

Herbarium vouchers for all species were deposited at Rocky Mountain Biological Laboratory (RMBL) herbarium (Appendix 1) and New York Botanical Garden William and Lynda Steere Herbarium. Due to a backlog of vouchers at NYBG, our specimens had not yet been processed at the time of publication. They will be able to be found using the collector’s and species’ names (R. Cohen) after they have been fully processed by NYBG.

All genomic and transcriptomic data will be available on NCBI, and we will provide the SRAs at the time of publication. Dataframes containing pollen tube growth rate data will be available on Dryad.

## Author Contributions

R.O.C and D.E. contributed equally to transcriptome study design. R.O.C. and J.H.W. contributed to pollen tube growth rate study design. R.O.C. collected all data and conducted all differential expression analyses. R.O.C., A.M., and S.M.C. conducted pistil staining and microscopy. R.O.C. and S.M.C. conducted pollen tube growth rate statistical analyses. R.O.C., D.E., A.M., and S.M.C, and J.H.W. contributed to the writing of the manuscript.

## Acknowledgements

We thank J. Jones for her assistance in RNA collection, and E. Leclair and E. Jedlicka for their work collecting samples in the field for pollen tube growth rate quantification. Thanks to Jennifer Reithel and Rick Williams their help onsite at RMBL. Special thanks to J. Coughlan for their feedback on the manuscript.

## Literature cited

Alves, C. M. L., A. K. Noyszewski, and A. G. Smith. 2019. Nicotiana tabacum pollen–pistil interactions show unexpected spatial and temporal differences in pollen tube growth among genotypes. Plant Reprod. 32:341–352.

Arceo-Gómez, G., R. L. Kaczorowski, C. Patel, and T.-L. Ashman. 2019. Interactive effects between donor and recipient species mediate fitness costs of heterospecific pollen receipt in a co-flowering community. Oecologia 189:1041–1047.

Ashman, T.-L., and G. Arceo-Gómez. 2013. Toward a predictive understanding of the fitness costs of heterospecific pollen receipt and its importance in co-flowering communities. Am. J. Bot. 100:1061–1070.

Baek, Y. S., P. A. Covey, J. J. Petersen, R. T. Chetelat, B. McClure, and P. A. Bedinger. 2015. Testing the SI × SC rule: Pollen–pistil interactions in interspecific crosses between members of the tomato clade (Solanum section Lycopersicon, Solanaceae). Am. J. Bot. 102:302–311.

Bedinger, P. A., A. K. Broz, A. Tovar-Mendez, and B. McClure. 2017. Pollen-Pistil Interactions and Their Role in Mate Selection. Plant Physiol. 173:79–90.

Bodenhofer, U., E. Bonatesta, C. Horejš-Kainrath, and S. Hochreiter. 2015. msa: an R package for multiple sequence alignment. Bioinformatics 31:3997–3999.

Briggs, H. M., L. M. Anderson, L. M. Atalla, A. M. Delva, E. K. Dobbs, and B. J. Brosi. 2016. Heterospecific pollen deposition in Delphinium barbeyi: linking stigmatic pollen loads to reproductive output in the field. Ann. Bot. 117:341–347.

Brosi, B. J., and H. M. Briggs. 2013. Single pollinator species losses reduce floral fidelity and plant reproductive function. Proc. Natl. Acad. Sci. 110:13044–13048. Proceedings of the National Academy of Sciences.

Broz, A. K., and P. A. Bedinger. 2021. Pollen-Pistil Interactions as Reproductive Barriers. Annu. Rev. Plant Biol. 72:615–639.

Cao, Y., X. Cui, Y. Yang, L. Pan, F. Yang, S. Li, D. Wu, Y. Ding, R. Chen, N. Wang, S. Liu, Z. Ji, Y. Zhao, Y. Chen, R. Sun, S. Xian, L. Yang, J. Hui, R. Li, T. Zhang, S. Dou, G. Song, X. Wei, Y. Yuan, X. Zhang, M. Chen, X. Sun, H.-M. Wu, A. Y. Cheung, and Q. Duan. 2025. Pan-family pollen signals control an interspecific stigma barrier across Brassicaceae species. Science 0:eady2347. American Association for the Advancement of Science.

Cheng, C.-Y., V. Krishnakumar, A. P. Chan, F. Thibaud-Nissen, S. Schobel, and C. D. Town. 2017. Araport11: a complete reannotation of the Arabidopsis thaliana reference genome. Plant J. 89:789–804.

Cohen, R. O., A. Chomentowska, L. Cai, and D. A. R. Eaton. 2025a. A reference genome and transcriptome of haustorial development in Pedicularis groenlandica reveal diverse trajectories of haustoria-associated gene evolution in parasitic plants. bioRxiv.

Cohen, R. O., A. Cisse, J. U. Jones, J. H. Williams, and D. A. R. Eaton. 2025b. Phylogeny does not predict the outcome of heterospecific pollen–pistil interactions in a species-rich alpine plant community. Am. J. Bot. n/a:e70004.

Coughlan, J. M., M. Wilson Brown, and J. H. Willis. 2020. Patterns of Hybrid Seed Inviability in the *Mimulus guttatus* sp. Complex Reveal a Potential Role of Parental Conflict in Reproductive Isolation. Curr. Biol. 30:83–93.e5.

Coyne, J. A., and H. A. Orr. 2004. Speciation. Oxford University Press, Oxford, New York.

de Nettancourt, D. 1997. Incompatibility in angiosperms. Sex. Plant Reprod. 10:185–199.

Devireddy, A. R., T. J. Tschaplinski, G. A. Tuskan, W. Muchero, and J.-G. Chen. 2021. Role of Reactive Oxygen Species and Hormones in Plant Responses to Temperature Changes. Int. J. Mol. Sci. 22:8843. Multidisciplinary Digital Publishing Institute.

Dobin, A., C. A. Davis, F. Schlesinger, J. Drenkow, C. Zaleski, S. Jha, P. Batut, M. Chaisson, and T. R. Gingeras. 2013. STAR: ultrafast universal RNA-seq aligner. Bioinforma. Oxf. Engl. 29:15–21.

Duan, Q., D. Kita, E. A. Johnson, M. Aggarwal, L. Gates, H.-M. Wu, and A. Y. Cheung. 2014. Reactive oxygen species mediate pollen tube rupture to release sperm for fertilization in Arabidopsis. Nat. Commun. 5:3129. Nature Publishing Group.

Eaton, D. A. R., C. B. Fenster, J. Hereford, S.-Q. Huang, and R. H. Ree. 2012. Floral diversity and community structure in Pedicularis (Orobanchaceae). Ecology 93:S182–S194.

Emms, D. M., and S. Kelly. 2019. OrthoFinder: phylogenetic orthology inference for comparative genomics. Genome Biol. 20:238.

Figueroa-Castro, D. M., and T. P. Holtsford. 2009. Post-pollination mechanisms in Nicotiana longiflora and N. plumbaginifolia: pollen tube growth rate, offspring paternity and hybridization. Sex. Plant Reprod. 22:187–196.

Garlovsky, M. D., and R. R. Snook. 2018. Persistent postmating, prezygotic reproductive isolation between populations. Ecol. Evol. 8:9062–9073.

Garlovsky, M. D., E. Whittington, T. Albrecht, H. Arenas-Castro, D. M. Castillo, G. L. Keais, E. L. Larson, L. C. Moyle, M. Plakke, R. Reifová, R. R. Snook, M. Ålund, and A. A.-T. Weber. 2024. Synthesis and Scope of the Role of Postmating Prezygotic Isolation in Speciation. Cold Spring Harb. Perspect. Biol. 16:a041429.

Gilman, I. S., J. J. Moreno-Villena, Z. R. Lewis, E. W. Goolsby, and E. J. Edwards. 2022. Gene co-expression reveals the modularity and integration of C4 and CAM in Portulaca. Plant Physiol. 189:735–753.

Hamlin, J. A. P., N. A. Sherman, and L. C. Moyle. 2017. Two Loci Contribute Epistastically to Heterospecific Pollen Rejection, a Postmating Isolating Barrier Between Species. G3 GenesGenomesGenetics 7:2151–2159.

Herrero, M., and J. I. Hormaza. 1996. Pistil strategies controlling pollen tube growth. Sex. Plant Reprod. 9:343–347.

Hersch, E. I., and B. A. Roy. 2007. CONTEXT-DEPENDENT POLLINATOR BEHAVIOR: AN EXPLANATION FOR PATTERNS OF HYBRIDIZATION AMONG THREE SPECIES OF INDIAN PAINTBRUSH. Evolution 61:111–124.

Hirose, T., A. Ujihara, H. Kitabayashi, and M. Minami. 1995. Pollen Tube Behavior Related to Self-incompatibility in Interspecific Crosses Of Fagopyrum. Jpn. J. Breed. 45:65–70. Japanese Society of Breeding.

Hogenboom, N. G., K. Mather, J. Heslop-Harrison, and D. Lewis. 1997. Incompatibility and incongruity: two different mechanisms for the non-functioning of intimate partner relationships. Proc. R. Soc. Lond. B Biol. Sci. 188:361–375. Royal Society.

Hormaza, J. I., and M. Herrero. 1994. Gametophytic competition and selection. Pp. 372–400 in E. G. Williams, A. E. Clarke, and R. B. Knox, eds. Genetic control of self-incompatibility and reproductive development in flowering plants. Springer Netherlands, Dordrecht.

Howard, D. J. 1999. Conspecific Sperm and Pollen Precedence and Speciation. Annu. Rev. Ecol. Evol. Syst. 30:109–132. Annual Reviews.

Jombart, T., F. Balloux, and S. Dray. 2010. adephylo: new tools for investigating the phylogenetic signal in biological traits. Bioinformatics 26:1907–1909.

Kaya, H., R. Nakajima, M. Iwano, M. M. Kanaoka, S. Kimura, S. Takeda, T. Kawarazaki, E. Senzaki, Y. Hamamura, T. Higashiyama, S. Takayama, M. Abe, and K. Kuchitsu. 2014. Ca2+-activated reactive oxygen species production by Arabidopsis RbohH and RbohJ is essential for proper pollen tube tip growth. Plant Cell 26:1069–1080.

Kodera, C., J. Just, M. Da Rocha, A. Larrieu, L. Riglet, J. Legrand, F. Rozier, T. Gaude, and I. Fobis-Loisy. 2021. The molecular signatures of compatible and incompatible pollination in Arabidopsis. BMC Genomics 22:268.

Kohn, J. R., and N. M. Waser. 1985. The Effect of Delphinium nelsonii Pollen on Seed Set in Ipomopsis aggregata, a Competitor for Hummingbird Pollination. Am. J. Bot. 72:1144–1148.

Krichevsky, A., S. V. Kozlovsky, G.-W. Tian, M.-H. Chen, A. Zaltsman, and V. Citovsky. 2007. How pollen tubes grow. Dev. Biol. 303:405–420.

Krueger, F. 2015. Trim Galore!: A wrapper around Cutadapt and FastQC to consistently apply adapter and quality trimming to FastQ files, with extra functionality for RRBS data.

Kumar, S., M. Suleski, J. M. Craig, A. E. Kasprowicz, M. Sanderford, M. Li, G. Stecher, and S. B. Hedges. 2022. TimeTree 5: An Expanded Resource for Species Divergence Times. Mol. Biol. Evol. 39:msac174.

Langfelder, P., and S. Horvath. 2008. WGCNA: an R package for weighted correlation network analysis. BMC Bioinformatics 9:559.

Larson, E. L., M. M. Brassil, J. Maslan, D. Juárez, F. Lilagan, H. Tipton, A. Schweitzer, J. Skillman, K. J. Monsen-Collar, and M. A. Peterson. 2019. The effects of heterospecific mating frequency on the strength of cryptic reproductive barriers. J. Evol. Biol. 32:900–912.

Liang, H., Z.-X. Ren, Z.-B. Tao, Y.-H. Zhao, P. Bernhardt, D.-Z. Li, and H. Wang. 2018. Impact of pre- and post-pollination barriers on pollen transfer and reproductive isolation among three sympatric Pedicularis (Orobanchaceae) species. Plant Biol. 20:662–673.

Liao, Y., G. K. Smyth, and W. Shi. 2014. featureCounts: an efficient general purpose program for assigning sequence reads to genomic features. Bioinforma. Oxf. Engl. 30:923–930.

Lin, S.-Y., P.-W. Chen, M.-H. Chuang, P. Juntawong, J. Bailey-Serres, and G.-Y. Jauh. 2014. Profiling of Translatomes of in Vivo–Grown Pollen Tubes Reveals Genes with Roles in Micropylar Guidance during Pollination in Arabidopsis. Plant Cell 26:602–618.

Liu, B., N. Boivin, D. Morse, and M. Cappadocia. 2012. A time course of GFP expression and mRNA stability in pollen tubes following compatible and incompatible pollinations in Solanum chacoense. Sex. Plant Reprod. 25:205–213.

Lobaton, J., R. Andrew, J. Duitama, L. Kirkland, S. Macfadyen, and R. Rader. 2021. Using RNA-seq to characterize pollen–stigma interactions for pollination studies. Sci. Rep. 11:6635. Nature Publishing Group.

Lora, J., J. I. Hormaza, and M. Herrero. 2016. The Diversity of the Pollen Tube Pathway in Plants: Toward an Increasing Control by the Sporophyte. Front. Plant Sci. 7:107.

Losada, J. M., and M. Herrero. 2014. Glycoprotein composition along the pistil of Malus x domestica and the modulation of pollen tube growth. BMC Plant Biol. 14:1.

Love, M. I., W. Huber, and S. Anders. 2014. Moderated estimation of fold change and dispersion for RNA-seq data with DESeq2. Genome Biol. 15:550.

Luu, D.-T., X. Qin, D. Morse, and M. Cappadocia. 2000. S-RNase uptake by compatible pollen tubes in gametophytic self-incompatibility. Nature 407:649–651. Nature Publishing Group.

Macior, L. W. 1995. Pollination Ecology of Pedicularis in the Teton Mountain Region. Plant Species Biol. 10:77–82.

Macior, L. W. 1983. The Pollination Dynamics of Sympatric Species of Pedicularis (scrophulariaceae). Am. J. Bot. 70:844–853.

Macior, L. W. 1970. The Pollination Ecology of Pedicularis in Colorado. Am. J. Bot. 57:716–728.

Mizuta, Y., and T. Higashiyama. 2018. Chemical signaling for pollen tube guidance at a glance. J. Cell Sci. 131:jcs208447.

Moreira-Hernández, J. I., and N. Muchhala. 2019. Importance of Pollinator-Mediated Interspecific Pollen Transfer for Angiosperm Evolution. Annu. Rev. Ecol. Evol. Syst. 50:191–217.

Nasrallah, J., and M. Nasrallah. 1993. Pollen-Stigma Signaling in the Sporophytic Self-Incompatibility Response. Plant Cell 5:1325–1335.

Nelson, T. C., A. M. Stathos, D. D. Vanderpool, F. R. Finseth, Y. Yuan, and L. Fishman. 2021. Ancient and recent introgression shape the evolutionary history of pollinator adaptation and speciation in a model monkeyflower radiation (Mimulus section Erythranthe). PLOS Genet. 17:e1009095. Public Library of Science.

Paradis, E., and K. Schliep. 2019. ape 5.0: an environment for modern phylogenetics and evolutionary analyses in R. Bioinforma. Oxf. Engl. 35:526–528.

Qin, Y., A. R. Leydon, A. Manziello, R. Pandey, D. Mount, S. Denic, B. Vasic, M. A. Johnson, and R. Palanivelu. 2009. Penetration of the Stigma and Style Elicits a Novel Transcriptome in Pollen Tubes, Pointing to Genes Critical for Growth in a Pistil. PLOS Genet. 5:e1000621. Public Library of Science.

Ramsey, J., H. D. Bradshaw JR., and D. W. Schemske. 2003. Components of Reproductive Isolation Between the Monkeyflowers Mimulus Lewisii and M. Cardinalis (phrymaceae). Evolution 57:1520–1534.

Richman, A. D., and J. R. Kohn. 2000. Evolutionary genetics of self-incompatibility in the Solanaceae. Pp. 169–179 in J. J. Doyle and B. S. Gaut, eds. Plant Molecular Evolution. Springer Netherlands, Dordrecht.

Rifkin, J. L., K. L. Ostevik, and M. D. Rausher. 2023. Complex cross-incompatibility in morning glories is consistent with a role for mating system in plant speciation. Evolution 77:1691–1703.

Riglet, L., F. Rozier, C. Kodera, S. Bovio, J. Sechet, I. Fobis-Loisy, and T. Gaude. 2020. KATANIN-dependent mechanical properties of the stigmatic cell wall mediate the pollen tube path in Arabidopsis. eLife 9:e57282. eLife Sciences Publications, Ltd.

Sanders, L. C., and E. M. Lord. 1989. Directed Movement of Latex Particles in the Gynoecia of Three Species of Flowering Plants. Science 243:1606–1608. American Association for the Advancement of Science.

Sanders, L. C., and E. M. Lord. 1992. The Extracellular Matrix in Pollen Tube Growth. Pp. 238–244 in E. Ottaviano, M. S. Gorla, D. L. Mulcahy, and G. B. Mulcahy, eds. Angiosperm Pollen and Ovules. Springer, New York, NY.

Schliep, K. P. 2011. phangorn: phylogenetic analysis in R. Bioinformatics 27:592–593.

Steinhorst, L., and J. Kudla. 2013. Calcium and Reactive Oxygen Species Rule the Waves of Signaling1. Plant Physiol. 163:471–485.

Streher, N. S., P. J. Bergamo, T.-L. Ashman, M. Wolowski, and M. Sazima. 2020. Effect of heterospecific pollen deposition on pollen tube growth depends on the phylogenetic relatedness between donor and recipient. AoB PLANTS 12:plaa016.

Swanson, W. J., and V. D. Vacquier. 2002. The rapid evolution of reproductive proteins. Nat. Rev. Genet. 3:137–144. Nature Publishing Group.

Swindell, W. R., M. Huebner, and A. P. Weber. 2007. Transcriptional profiling of Arabidopsis heat shock proteins and transcription factors reveals extensive overlap between heat and non-heat stress response pathways. BMC Genomics 8:125.

Takayama, S., and A. Isogai. 2005. Self-incompatibility in plants. Annu. Rev. Plant Biol. 56:467–489.

Taylor, L. P., and P. K. Hepler. 1997. POLLEN GERMINATION AND TUBE GROWTH. Annu. Rev. Plant Biol. 48:461–491. Annual Reviews.

Tikhonov, G., Ø. H. Opedal, N. Abrego, A. Lehikoinen, M. M. J. de Jonge, J. Oksanen, and O. Ovaskainen. 2020. Joint species distribution modelling with the r-package Hmsc. Methods Ecol. Evol. 11:442–447.

Ting, J. J., G. C. Woodruff, G. Leung, N.-R. Shin, A. D. Cutter, and E. S. Haag. 2014. Intense Sperm-Mediated Sexual Conflict Promotes Reproductive Isolation in Caenorhabditis Nematodes. PLOS Biol. 12:e1001915. Public Library of Science.

Tong, Z.-Y., and S.-Q. Huang. 2016. Pre- and post-pollination interaction between six co-flowering Pedicularis species via heterospecific pollen transfer. New Phytol. 211:1452–1461.

Visser, T. 1981. Pollen and pollination experiments. IV. ‘Mentor pollen’ and ‘pioneer pollen’ techniques regarding incompatibility and incongruity in apple and pear. Euphytica 30:363–369.

Visser, T. 1983. THE ROLE OF PIONEER POLLEN IN COMPATIBLE AND INCOMPATIBLE POLLINATIONS OF APPLE AND PEAR. Acta Hortic. 51–58.

Wu, J., Y. Qin, and J. Zhao. 2008. Pollen tube growth is affected by exogenous hormones and correlated with hormone changes in styles in Torenia fournieri L. Plant Growth Regul. 55:137–148.

Wu, T., E. Hu, S. Xu, M. Chen, P. Guo, Z. Dai, T. Feng, L. Zhou, W. Tang, L. Zhan, X. Fu, S. Liu, X. Bo, and G. Yu. 2021. clusterProfiler 4.0: A universal enrichment tool for interpreting omics data. The Innovation 2. Elsevier.

Ye, J., S. McGinnis, and T. L. Madden. 2006. BLAST: improvements for better sequence analysis. Nucleic Acids Res. 34:W6–W9.

