## Supplemental Figures for "From recognition to neglect: Molecular and physiological responses to heterospecific pollen decay with evolutionary distance"

*P. groenlandica* x *P. groenlandica*, Individual #93, Timepoint 1

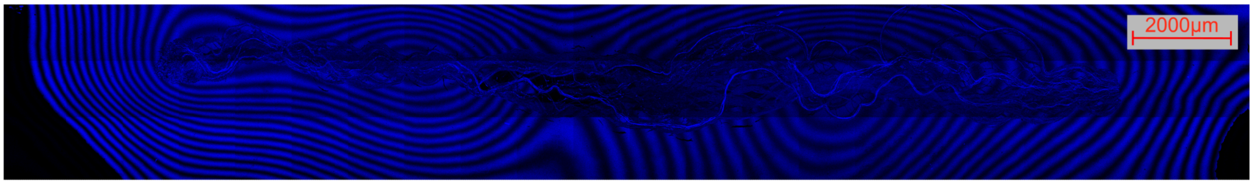


*P. groenlandica* x *P. groenlandica*, Individual #1, Timepoint 2

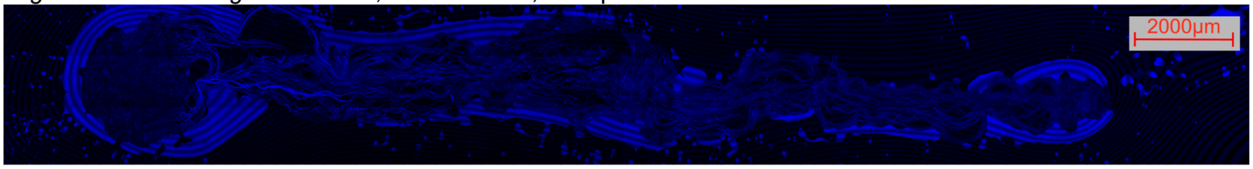


*P. procera* x *P. procera*, Individual #23, Timepoint 2

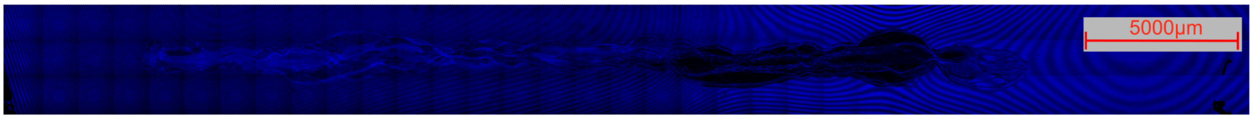


*C. sulphurea* x *C. sulphurea*, Individual #22, Timepoint 2

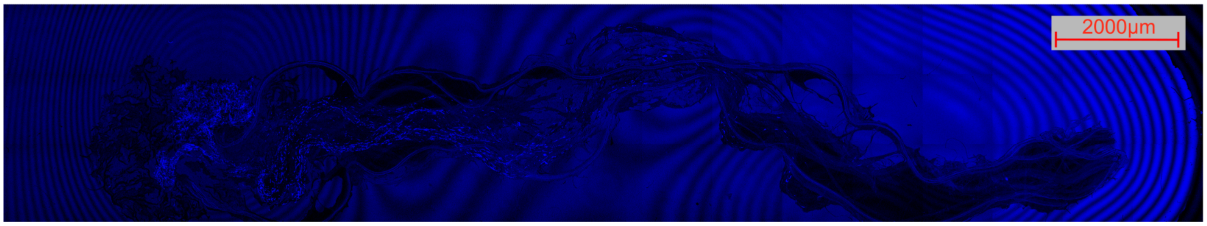


**Fig. S1.** Examples of fluorescence microscopy images showing aniline blue stained pistils from which pollen tube growth rates were measured. Pollen tubes were dim in some images, but could be seen by using zoom and increasing contrast.

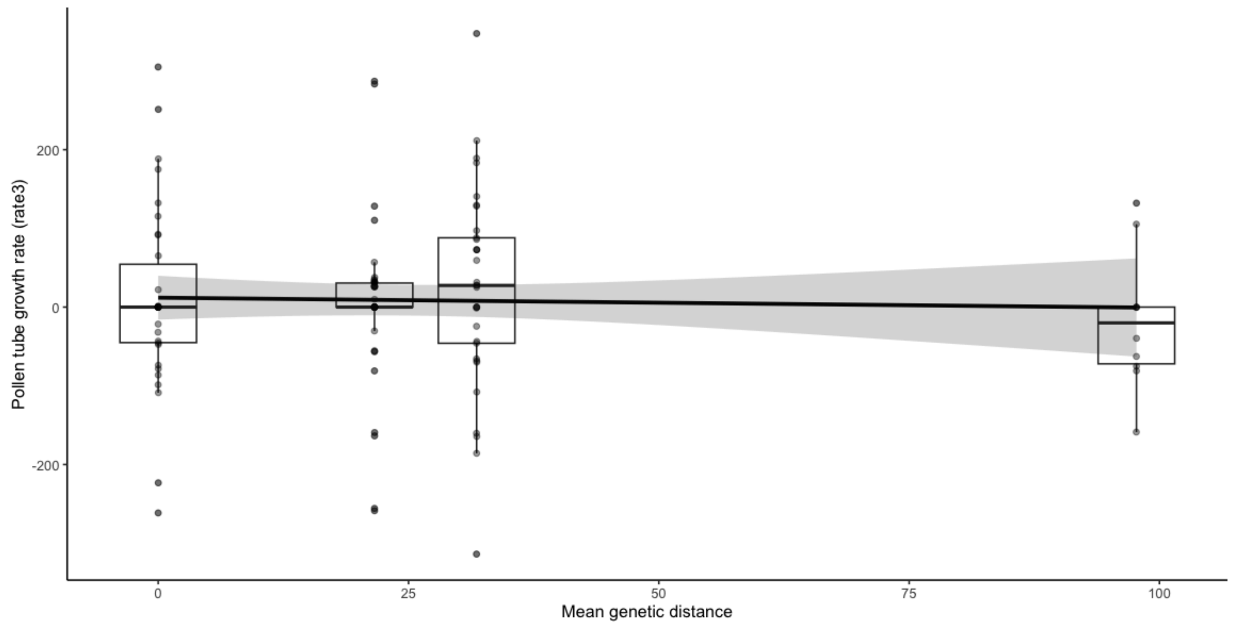


**FIg. S2.** Gaussian distributed generalized linear model fit to rate data calculated by subtracting length at T2-T1. We were only able to pair 110 datapoints total, and some T2 lengths were shorter than T1 lengths, resulting in negative rate values.

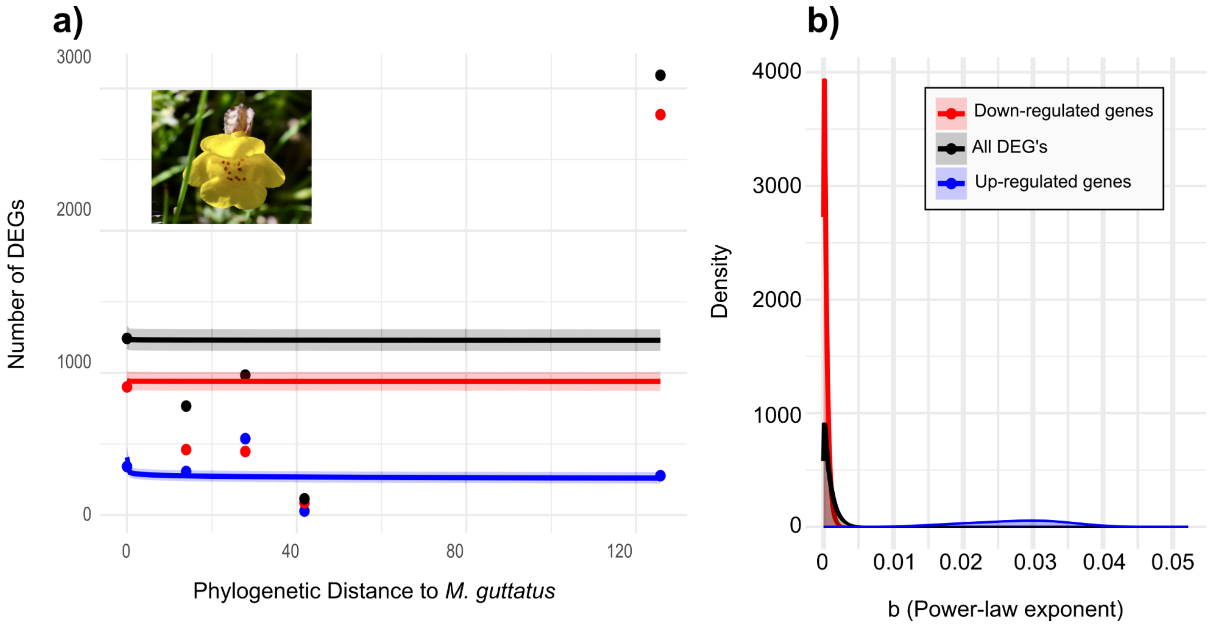


**Fig. S3.** Negative power law model fit to *M. guttatus* cross DEG data when the outlier *Mimulus guttatus* X *Prunella vulgaris* is included. **a)** Although there is decay in the number of DEGs according to phylogenetic distance until the *M. guttatus* X *P. groenlandica* cross (*see* **Fig. 2**), there is no relationship between between phylogenetic distance and number of DEGs when *M. guttatus* X *P. vulgaris* is included. The *M. guttatus* X *P. vulgaris* had the highest number of DEGs of any cross in the study, including conspecific crosses. The majority of these DEGs were down-regulated (shown in red), as compared to up-regulated (blue) and all DEGs (black). There is minor negative relationship between number of DEGs and phylogentic distance for only the up-regulated genes, as shown in the **b)** posterior density curve for the b (power-law exponent) parameter

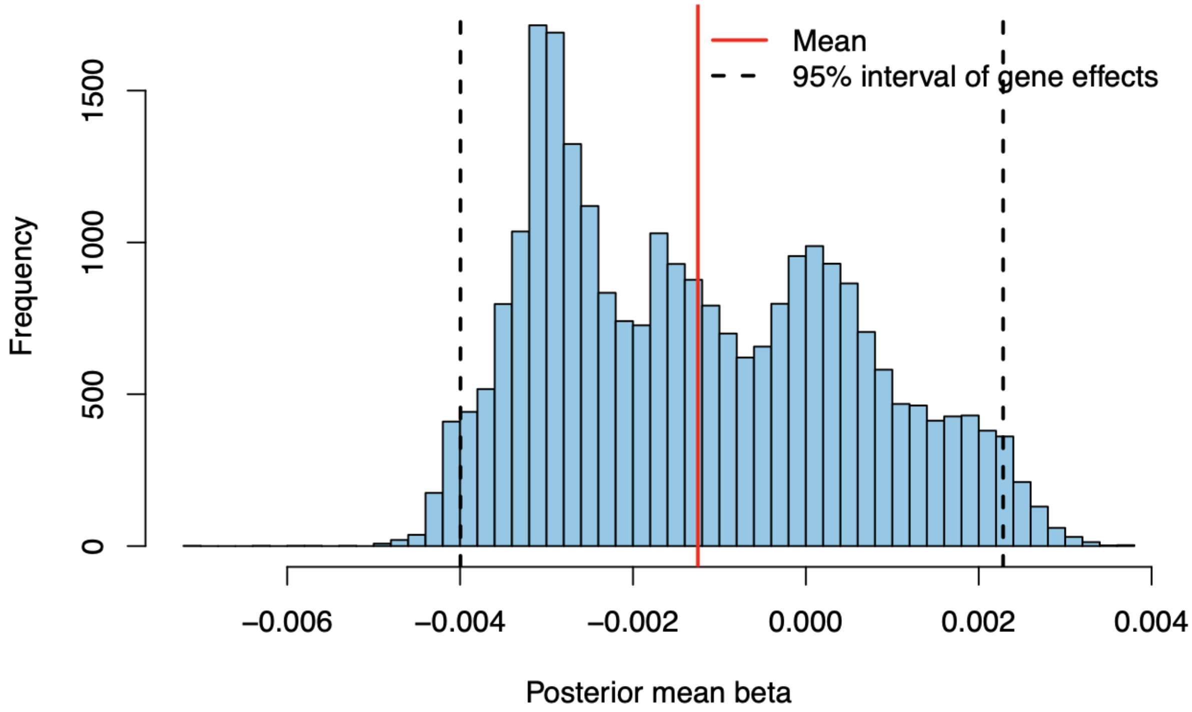


**Fig. S4.** Histogram showing the distribution of mean beta estimates per gene from the hierarchical model. This model was fit only to the differential expression data from crosses to *P. groenlandica*. Here, beta describes the relationship between phylogenetic distance and per gene variance stabilized expression. The majority (~78%) of beta means were negative, indicating that many genes were less likely to be expressed as the phylogenetic distance between *P. groenlandica* and the pollen donor species increased.
